# AI.zymes – A modular platform for evolutionary enzyme design

**DOI:** 10.1101/2025.01.18.633707

**Authors:** Lucas P. Merlicek, Jannik Neumann, Abbie Lear, Vivian Degiorgi, Moor de Waal, Tudor-Stefan Cotet, Adrian J. Mulholland, H. Adrian Bunzel

## Abstract

The ability to create new-to-nature enzymes would substantially advance bioengineering, medicine, and the chemical industry. Despite recent breakthroughs in protein design and structure prediction, designing biocatalysts with activities rivaling those of natural enzymes remains challenging. Here, we present AI.zymes, a modular platform integrating cutting-edge protein engineering algorithms within an evolutionary framework. By combining programs such as Rosetta, ESMFold, ProteinMPNN, and FieldTools in iterative rounds of design and selection, AI.zymes optimizes a broad range of catalytically relevant properties, such as transition state affinity and protein stability. Notably, AI.zymes can also improve properties that are not targeted by the employed design algorithms. For instance, AI.zymes enhanced electrostatic catalysis by iteratively selecting variants with stronger catalytic electric fields. AI.zymes was benchmarked by improving the promiscuous Kemp eliminase activity of ketosteroid isomerase, yielding a 7.7-fold increase in activity after zero-shot experimental validation of only 7 variants. Due to its modularity, AI.zymes can readily incorporate emerging design algorithms, paving the way for a unifying framework for enzyme design.

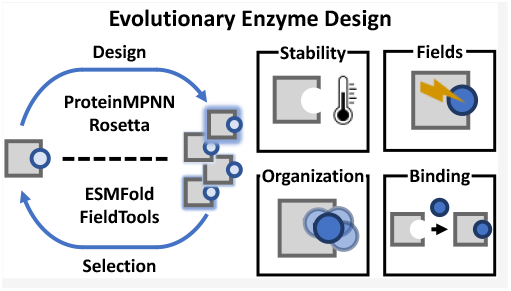

---

Enzymes are nature’s catalysts, facilitating chemical reactions with remarkable specificity and efficiency. The ability to create tailor-made enzymes holds immense potential for biotechnology, medicine, and the chemical industry.^1,2^ However, the vastness of sequence space and the complex relationship between structure and function make enzyme design and engineering a significant challenge. Directed evolution offers a powerful approach to navigate these obstacles in iterative cycles of mutagenesis and selection to explore sequence space and optimize enzyme function.^3–5^ Yet, laboratory evolution is time-consuming and struggles to access distant regions of the fitness landscape.

Here, we present AI.zymes, an evolutionary design platform for optimizing enzymes in iterative rounds of design and selection *in silico* (Fig. 1a). Unlike non-evolutionary design algorithms based on a fixed input sequence,^6–10^ evolutionary design involves iteratively designing variants and selecting the most promising hits as templates for the next round of design.^11,12^ Varying the design input structure allows efficient exploration of sequence space towards regions inaccessible within a single design cycle. While computational evolution has proven effective in designing protein and nucleic acid binders,^11,12^ challenges in accurately modeling and scoring biocatalysts have, thus far, constrained enzyme engineering by evolutionary design.

**Fig. 1.**
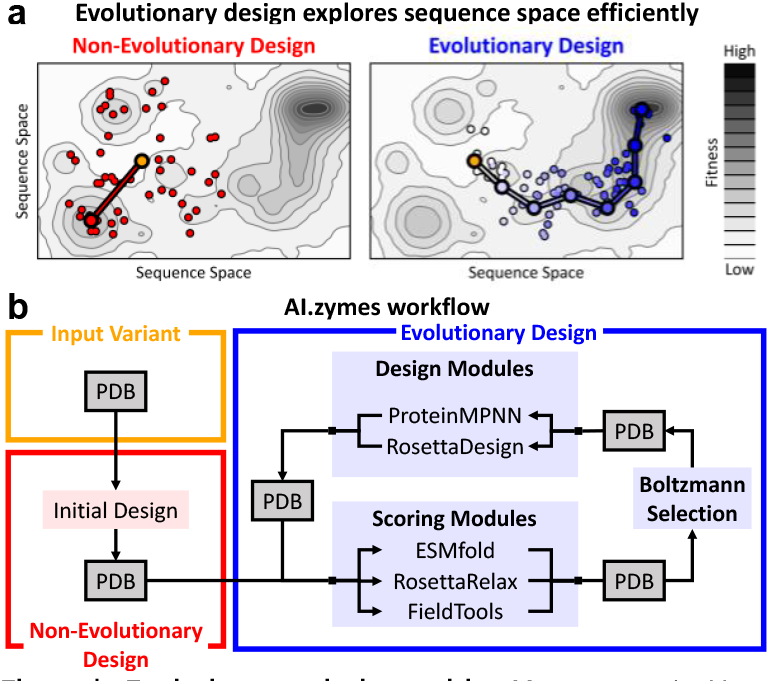
Evolutionary design with AI.zymes. **a)** Non-evolutionary design (red) starts from a fixed input variant (yellow), which limits diversity. In contrast, evolutionary design (blue) explores sequence space more efficiently. **b)** Modularity in AI.zymes is achieved by linking standalone algorithms through their output structures (PDB, grey).

Recent protein design and structure prediction breakthroughs have substantially boosted our enzyme engineering abilities. Tools such as AlphaFold3 and ESMFold accurately predict protein structures to inform engineering,^13,14^ and generative models such as ProteinMPNN and RFdiffusion reliably produce *de novo* proteins.^8,9^ These advances recently culminated in the *de novo* design of several enzymes by creating novel protein scaffolds around active-site models.^15,16^

While impressive, even the best designer enzymes generally do not achieve the catalytic power of natural enzymes.^15,16^ Common protein design algorithms aim at maximizing protein stability.^6–10^ Due to a lack of suitable tools, enzyme design thus often focuses on active-site shape complementarity while overlooking critical electrostatic and dynamic factors governing enzyme activity.^3,19,20,23^ Biocatalytically important effects, such as electrostatic catalysis^17,18^ and conformational dynamics,^3,19,20^ can usually only be targeted by filtering variants after design.^21,22^ This methodological gap probably contributes to the low *k*cat values of *de novo* enzymes,^15,16^ underscoring the need for a holistic multi-objective framework to optimize a range of biocatalytically relevant properties during design.

Here, we present AI.zymes, a modular platform for evolutionary design that integrates multiple design methods such as Rosetta, ESMFold, and ProteinMPNN (Fig. 1b). AI.zymes was benchmarked on the promiscuous Kemp eliminase activity of ketosteroid isomerase. Optimization of transition-state recognition, protein stability, and catalytic electric fields resulted in a 7.7-fold activity improvement in zero-shot validation. AI.zymes provides a general framework for enzyme design by modularly integrating established design algorithms in a multi-objective optimization pipeline.

## RESULTS

### How AI.zymes works

Similar to non-evolutionary design approaches, AI.zymes starts with generating an initial pool of variants based on an input structure. Subsequently, the best variants from the growing pool are selected for redesign and scored with various computational enzyme activity metrics (Fig. S1).

During selection, AI.zymes first eliminates variants that do not match the target geometry between the chemical transition state and catalytic residues.^24^ Then, multi-objective Boltzmann selection is performed based on scores describing the ligand-protein interaction strength (interface score),^25^ protein stability (stability score), and electrostatic catalysis (electric fields,^23^ Fig. 2a). AI.zymes runs evolution in a forward-thinking manner, where the scores of the direct descendants of a variant affect that variant’s likelihood to be redesigned. This is achieved by selecting based on a metric called “potential” corresponding to the average score of each variant and its first-generation descendants. To achieve multi-objective optimization, Boltzmann selection is based on the average of all relevant standardized potentials. Finally, AI.zymes gradually adjusts the Boltzmann temperature during selection to transition from an exploration phase where input structures are chosen more randomly to an exploitation phase that is highly selective for the most promising variants.

**Fig. 2.**
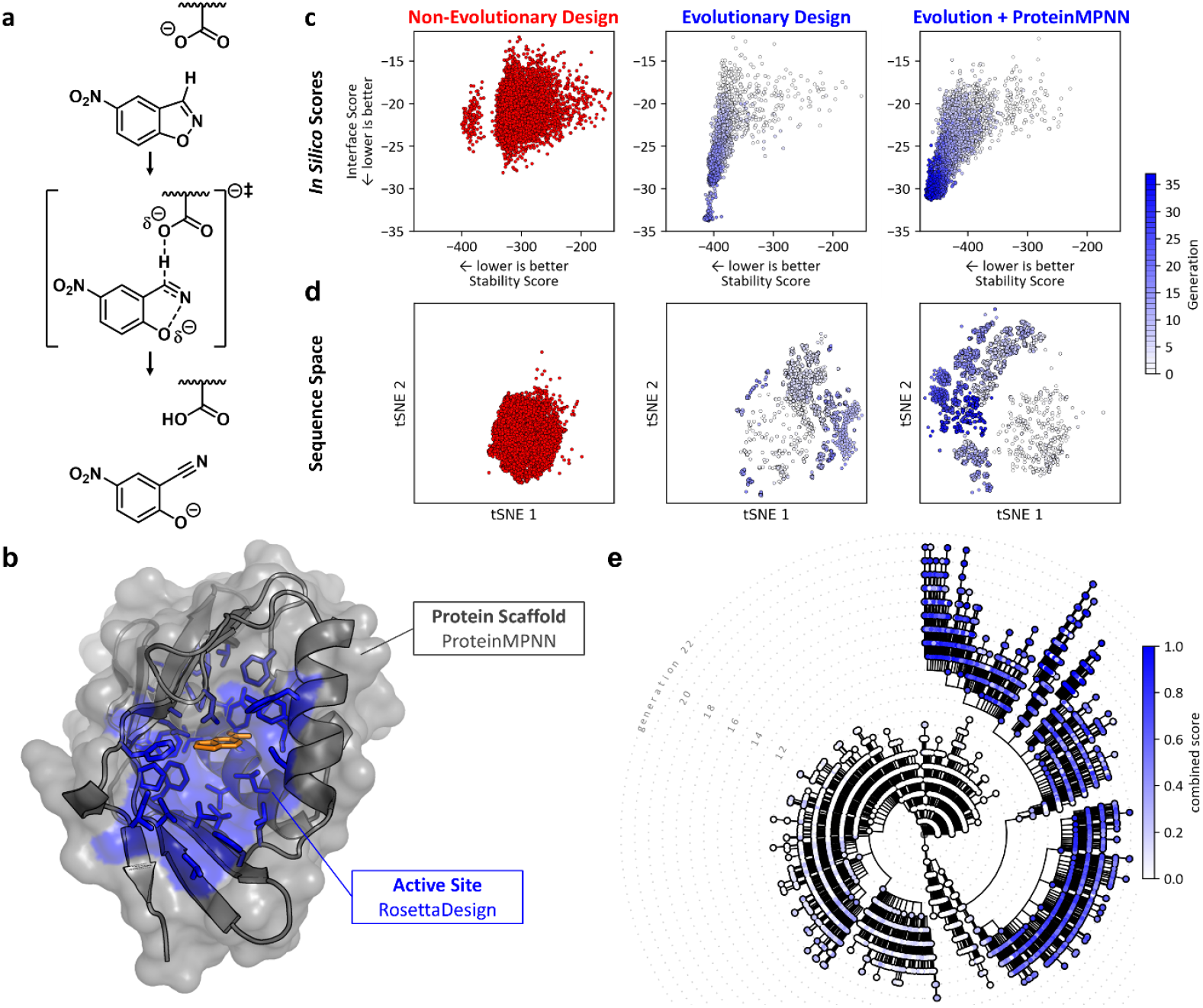
Evolutionary design outperforms non-evolutionary design. **a)** AI.zymes was benchmarked by improving the Kemp eliminase activity of KSI. **b)** The active site of KSI was targeted by RosettaDesign (blue) and the protein scaffold was redesigned with ProteinMPNN (grey; transition state: orange; model based on PDB ID 1OHP)^26^. **c+d)** Compared to non-evolutionary design (red), evolutionary design (blue) achieved **(c)** higher *in silico* scores and **(d)** explored a broader sequence space (tSNE analysis of ESM2-embedded sequences). **e)** Phylogenetic tree tracing ancestors during computational evolution. The combined score represents the standardized mean of all scores targeted during design.

After Boltzmann selection, variants are subjected to structure prediction with ESMFold^13^ and relaxation using RosettaRelax^7,27^ to evaluate conformational stability and protein backbone shifts. Once relaxed, the resulting structures undergo redesign. In the current implementation of AI.zymes, active-site residues are optimized with RosettaDesign,^8,9^ whereas the protein scaffold is targeted using ProteinMPNN^9^ (Fig. 2b). This division leverages the strengths of each algorithm: Rosetta readily accounts for the bound chemical transition state, while ProteinMPNN excels at enhancing protein stability. To combine the design methods, each is applied with a user-defined probability during redesign (RosettaDesign: 70%; ProteinMPNN: 30%). Modularity in AI.zymes is based on design functions that use PDB files as input and output (Fig. 2b). For ProteinMPNN, this is achieved by linking the designed sequence to structure prediction and relaxation. This modularity facilitates the future integration of additional design methods and ensures the continued applicability of AI.zymes in the rapidly evolving field of computational protein design.

To explore sequence space gradually during evolutionary design, the mutational load during each design step must be limited (Tab. S1). Too many mutations per step would obliterate any evolutionary information embedded in the sequence, while too few restrict the overall evolutionary efficiency. To limit the mutational load, a weight was added to RosettaDesign that penalizes mutations relative to the input structure. For ProteinMPNN, this was similarly achieved by adding a bias towards the sequence of the input structure. Thus, mutation rate, Boltzmann selection temperature, and design method are crucial parameters to fine-tune the performance of AI.zymes.

### Evolutionary design outperforms non-evolutionary design

The promiscuous Kemp eliminase activity of ketosteroid isomerase (KSI) was selected as a target to benchmark AI.zymes.^28^ The Kemp elimination (the base-catalyzed C-H deprotonation of benzisoxazoles) is an established benchmark for *de novo* enzymes.^3,29–31^ KSI is a dimeric enzyme that promiscuously catalyzes the Kemp elimination of 5-nitrobenzisoxazole. The Kemp eliminase activity of *Comamonas testosterone* KSI has previously been improved by introducing a D38N mutation, removing one of the two active-site aspartates that could act as a catalytic base.^28^ Initial characterization confirmed that KSI D38N, referred to as WT hereafter, exhibits measurable Kemp eliminase activity, although size-exclusion chromatography revealed that the protein formed an oligomeric mixture(Fig. 4b). Therefore, AI.zymes was tasked with both enhancing activity and creating a thermostable and monomeric enzyme.

To benchmark AI.zymes, initial design runs were performed solely using RosettaDesign to optimize the active-site residues and ESMFold and RosettaRelax for structure prediction (Fig. 2b). After fine-tuning the Boltzmann selection temperature and mutation rate (Tab. S1), computational evolution outperformed non-evolutionary design in terms of interface and stability score (Fig. 1c), while exploring substantially broader sequence space. Integration of ProteinMPNN, which excels in creating thermostable proteins,^9^ into AI.zymes further improved the designs by increasing the stability score and expanding the explored sequence space. AI.zymes readily tracks the evolutionary design history. This functionality enabled the construction of a phylogenetic tree that maps ancestral variants to their descendants, providing a clear visualization of the incremental improvements during design (Fig. 2e).

### Evolutionary electric field design

Catalytic electric fields are critical in stabilizing transition states and promoting enzyme efficiency (Fig. 3a).^17,18^ Despite a few notable successes,^32,33^ the design of catalytic electric fields has been challenging. Furthermore, the limited *k*cat values of recent designer enzymes^3,15,16^ can probably be attributed partly to the neglect of electrostatic catalysis. To address this gap in current design methodologies, we integrated FieldTools, which we recently developed to assess electric fields from MD trajectories,^23^ into AI.zymes (Fig. S1).

**Fig. 3.**
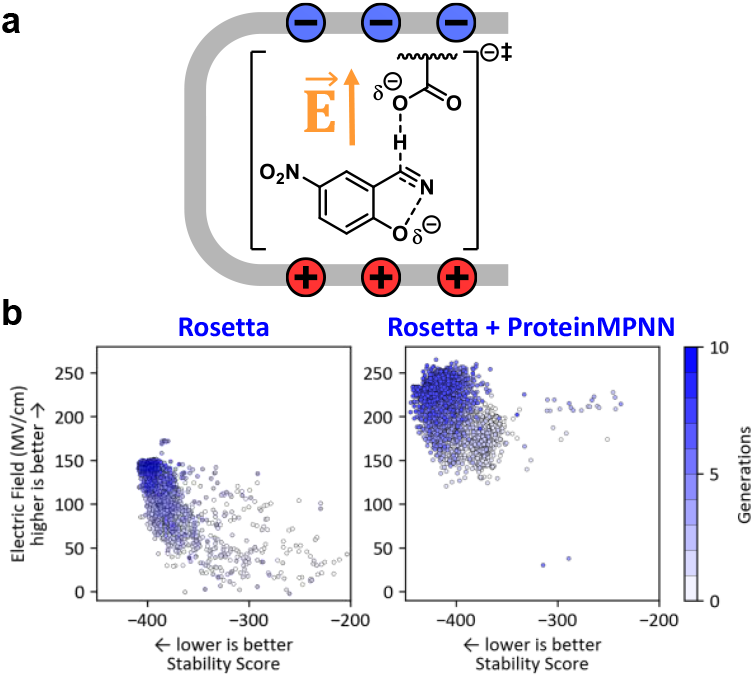
Designing catalytic electric fields. **a)** Electric fields (E, orange) quantify electrostatic catalysis. **b)** Design runs with ProteinMPNN and RosettaDesign result in higher catalytic electric fields than those using only RosettaDesign.

**Fig. 4.**
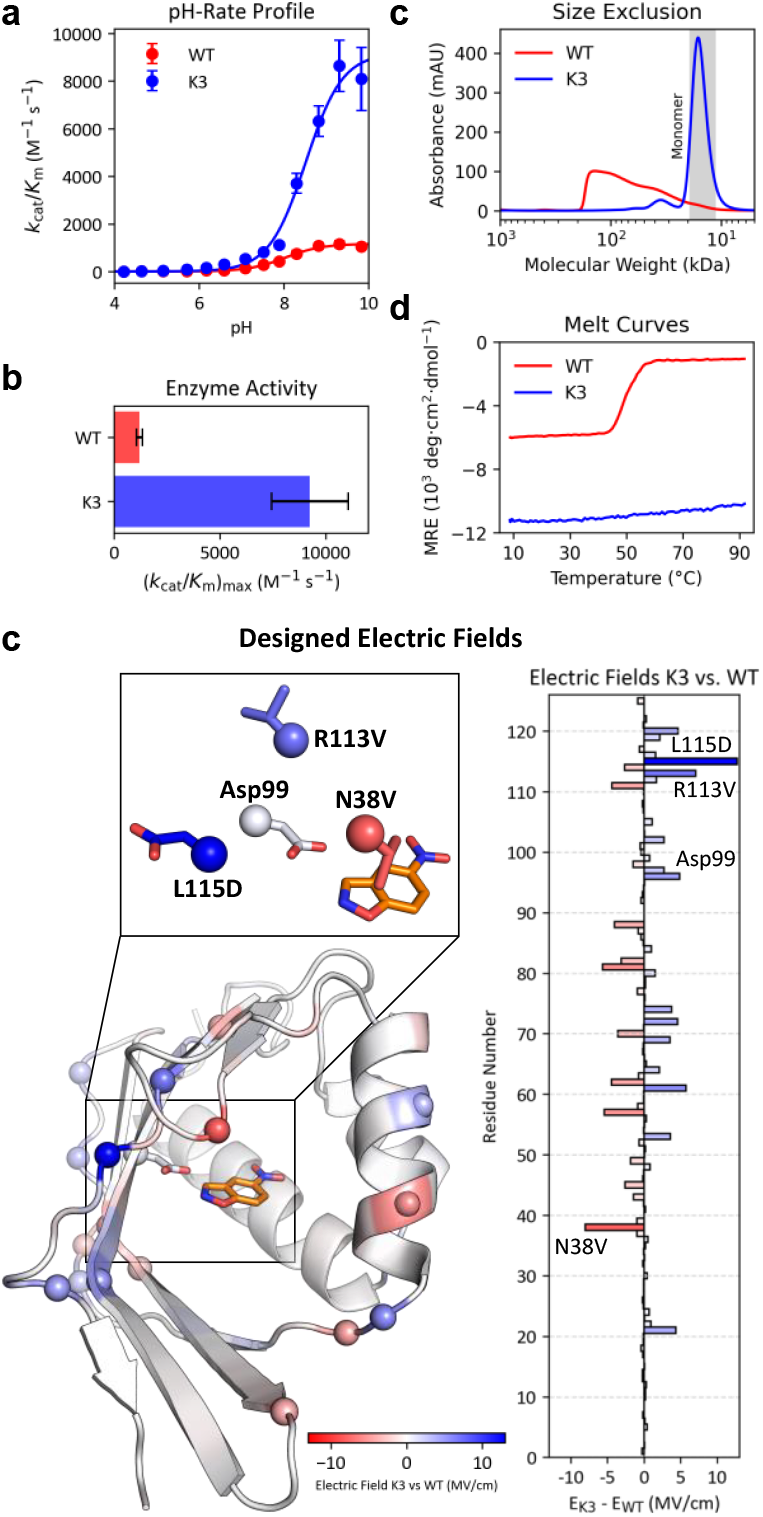
Evolutionary design improved enzyme activity. **a+b)** pH-Rate profiles reveal that K3 is up to 7.7 times more active than WT. Error bars indicate the standard error of 4 replicates. c) K3 is monomeric, whereas WT forms an oligomeric mixture. d) While WT unfolds at 49.8°C, K3 is stable up to 95°C. **e)** Per-residue comparison of the electric field in K3 versus WT reveals the origins of the increased field (increased field: blue; decreased field: red). **f)** Snapshots of the active sites of K3 (red) and WT (blue) from MD simulations indicate the strengthening of an H-bond from Tyr55 to the nitro group of the chemical transition state contributes to activity enhancement.

Notably, neither RosettaDesign nor ProteinMPNN natively support the targeted design of catalytic electric fields. In AI.zymes, electric field optimization is achieved by combining protein design agnostic to field effects with Boltzmann selection enriching variants with strong electric fields (Fig. 3b). The best fields were achieved in design runs that combined Rosetta and ProteinMPNN, as opposed to those relying solely on Rosetta. Thus, remote mutations in the protein scaffold played a crucial role in enhancing the fields. These results demonstrate that reliable electric field optimization—a significant advancement in enzyme design—can be achieved using methods not explicitly tailored for field effects.

### Molecular dynamics screening

In our previous work on the design of photoenzymes, we observed that short (10 ns) MD simulations are suitable for assessing the conformational stability of an enzyme-bound ligand.^22^ Here, the top 96 designs were screened using 10 ns molecular dynamics simulations to narrow the number of designs for experimental testing (Fig. S2).^34–42^ Variants were filtered by (1) ligand flexibility, (2) ligand-base distance, (3) electric field along the scissile C-H bond, (4) preorganization of the apo compared to the holo structure, and (5) active-site hydration. MD filtering identified 23 variants, of which 7 were chosen for experimental testing after visual inspection of the active-site geometry.

### Experimental Validation

Experimental validation is crucial to the design of functional proteins. Experimental activity assays at the highest possible substrate concentration (1.5 mM) revealed that 1 out of 7 tested variants was more active than WT, while the others had similar or lower activity (Fig. 4a+b and Tab. S3).^43^ pH-rate profiles of the improved variant, dubbed K3, revealed that its maximum *k*cat/*K*M was improved by 7.7-fold compared to WT, which was accompanied by a p*K*A shift of the general base from 8.1 to 8.4. Thus, at pH 7.0, the *k*cat/*K*m of K3 improved by 2.0-fold over WT, as shown by full Michaelis-Menten kinetics (Fig. S3c). Of 139 residues, 63 to 77 were mutated in the tested designs. Despite the high mutational load, only 7 designs had to be tested to find an improved variant. The experimental effort necessary to identify an improved design contrasts with typical computational design or machine-learning-assisted enzyme evolution projects that often require extensive experimental screening.

**Table 1.** Kinetics of WT and K3.

| variant | $(k_{cat}/K_M)_{max}$ ($M^{-1} s^{-1}$)<sup>a</sup> | $pK_A$ |
| --- | --- | --- |
| WT | $1200 \pm 200$ | $8.1 \pm 0.1$ |
| K3 | $9200 \pm 1800$ | $8.4 \pm 0.1$ |

<sup>a</sup>Maximum $k_{cat}/K_M$ determined from pH-rate profiles.

Size-exclusion chromatography revealed that the redesigned variants were predominantly monomeric, resolving the oligomerization issues found for the WT scaffold (Fig. 4c and Fig. S3b). Comparison of the pTM and iPTM scores from AlphaFold3 predictions for a range of WT and K3 oligomeric states supports that K3 is expected to form a stable monomer, while various oligomerization states were predicted for WT with high confidence (Fig. S4). Circular dichroism (CD) melting curves showed that K3 is highly thermostable and does not unfold up to 95°C, whereas WT had a melting temperature of 49.8°C. Interestingly, K3 was more stable and exhibited a stronger CD signal (Fig. S3d), which supports that multi-objective optimization can improve activity and afford a well-behaved and thermostable catalyst.

### Electric Field Optimization

Inspection of the mutations between WT and K3 revealed that the improvements in the total electric field arose from numerous small contributions (Fig. 4c). For instance, L115D introduced a negative charge that aids deprotonation, whereas R113V removes an anticatalytic positive charge in the WT. Interestingly, AI.zymes also introduced mutations such as N38V that negatively impact the electric field but enhance substrate binding by increasing the binding-site hydrophobicity. Notably, AI.zymes optimized the electric fields without relying on design tools tailored to design electrostatically beneficial mutations. This suggests that AI.zymes can optimize any aspect of enzyme function—even those that cannot be directly targeted by design—provided the desired trait can be scored and used in Boltzmann selection.

To probe how protein dynamics affect electrostatics, we analyzed K3 by MD simulations. These analyses revealed an increase in the electric field of 19 MV/cm compared to WT. The improvement was slightly lower than that calculated from the static structures (23 MV/cm), underscoring the importance of dynamics. In fact, repeated scoring of WT and K3 through ESMfold and RosettaRelax reproduced the score fluctuations seen in MD (Fig. S5). Beyond the electric fields, MD simulations highlighted other improved structural features in K3, including better ligand-base alignment and a strengthened hydrogen bond between Tyr55 and the nitro group of the chemical transition state (Fig. 4f). While these findings validate the results obtained from static structures, they also suggest that including the effects of dynamics on all calculated scores could further improve the design outcome.

## CONCLUSION

In this work, we developed AI.zymes, a modular framework that advances enzyme design through evolutionary multi-objective optimization. Using KSI as a model system, AI.zymes successfully improved the promiscuous Kemp eliminase activity by optimizing transition-state recognition, catalytic geometry, and electric fields while maintaining high protein stability. Integrating a range of cutting-edge computational tools, including RosettaDesign, ProteinMPNN, and ESMFold, enabled efficient exploration of the enzyme fitness landscape, which resulted in a 7.7-fold activity improvement after zero-shot experimental validation of just 7 variants.

A key advantage of evolutionary design is its ability to target mutagenesis within functional sequence space, which increases design efficiency compared to non-evolutionary methods. Increased efficiency is valuable for both large-scale design projects generating millions of designs and small-scale efforts constrained by limited computational resources. Boltzmann selection further enhances efficiency by operating on the entire design pool rather than using rounds-based approaches like experimental directed evolution. This approach maximizes resource utilization by continuously running a fixed number of designs without waiting for one round to finish before starting the next. Additionally, AI.zymes capability for multi-objective optimization—even for traits not directly addressed by the design algorithms, such as electrostatic catalysis—further improves functionality without increasing computational costs.

## OUTLOOK

Creating enzymes remains one of the major challenges in computational protein design. While recent design campaigns succeeded in generating some activity by creating protein scaffolds around active-site models,^15,16^ designing highly efficient enzymes will probably require extending beyond the design of defined protein-transition state complexes. Protein dynamics and electrostatic catalysis should be included during the design step – not as filters after the design – to more efficiently account for catalytically relevant conformational sampling and long-range effects within the protein scaffold.^3,17,19,20,32,33^ -

The modularity of AI.zymes renders it an ideal platform to advance enzyme design further. For instance, algorithms such as LigandMPNN, RFdiffusionAA, and AlphaFold3 that account for non-protein ligands could be integrated to model ligand-bound structures with increased accuracy.^14,44,45^ In addition, methods to probe protein dynamics, such as MD simulations or deep-learning tools (e.g., ChemNet^46^), could be included to enhance design. Our MD screening narrowed the range of design hits from 96 to 7 variants and suggests that short simulations (≤10 ns) are sufficient to eliminate designs with non-productive conformations. In total, 10^4^ designs were made during the design of K3, a number for which MD simulations informing Boltzmann selection could become computationally feasible.

AI.zymes succeeded in including catalytic electric fields in design, focusing on optimizing the electric field strength along a target bond. In the future, AI.zymes may furthermore be tasked with accounting for the electric field reorganization along the reaction coordinate and incorporating field effects across the entire transition state. A more refined description of electrostatic catalysis could help design more efficient designer enzymes.

Beyond optimizing enzyme electrostatics and dynamics, *de novo* design algorithms such as RosettaMatch, Combs, and Riff-Diff,^15,47^ could readily interface with AI.zymes to create enzyme activity from scratch. Moreover, active training of machine learning models during the design process could help guide exploration toward more active regions of the fitness landscape. Similar strategies have been applied in machine-learning-assisted directed evolution for experimental enzyme optimization^48^ and in training deep-learning models on sequence databases to develop new biocatalysts.^49–52^

Optimizing enzymes by targeting diverse properties such as protein stability, steric preorganization, and electrostatic catalysis is challenging for any single design algorithm. AI.zymes addresses this challenge with a modular platform integrating cutting-edge computational tools into a streamlined pipeline. Built on an evolutionary framework, AI.zymes combines the strengths of diverse methodologies in a multi-objective approach toward a unified algorithm for more effective enzyme design.

## AUTHOR CONTRIBUTIONS

HAB, LM, JN, and AL developed the code with support from TSC and VD. LM, VD, and MdW recorded the experimental data. HAB and LM wrote the manuscript with input from all co-authors.

## ACKNOWLEDGMENTS

We thank Basile Wicky for his helpful comments on our manuscript. HAB thanks the Swiss National Science Foundation for funding (Ambizione, PZ00P3_208691). This project has received funding from the European Research Council (ERC) under the European Horizon 2020 research and innovation program (PREDACTED Advanced Grant Agreement no. 101021207) to AJM. This work was conducted using the computational facilities of the Department of Biosystems Science and Engineering, ETH Zurich, and the Advanced Computing Research Centre, University of Bristol, http://www.bris.ac.uk/acrc/.

## DATA AVAILABILITY

Upon publication of the peer-reviewed manuscript, AI.zymes will be made available on GitHub. All data and methods required to reproduce this work will be made available via the ETH Research Collection and the supplementary information.

