## Supplementary Information for "AI.zymes – A modular platform for evolutionary enzyme design"

#### Contents

1. AI.zymes 3
1.1. Basic concepts 3
1.1.1. Evolutionary "rounds" in AI.zymes. 3
1.1.2. Scoring metrics. 3
1.2. System startup 4
1.3. Controller 5
1.4. Boltzmann selection. 5
1.4.1. Database 6
1.5. Available Modules in AI.zymes. 6
1.5.1. Design with RosettaDesign. 7
1.5.2. Design using ProteinMPNN. 7
1.5.3. Structure prediction with ESMfold and RosettaRelax. 8
1.5.4. Electric field calculations. 8
2. Molecular dynamics simulations 9
3. Experimental Methods 11
3.1. Protein production and purification 11
3.2. Enzyme kinetics 11
3.3. Circular dichroism 12
4. Supplementary Tables 13
Tab. S1 | Hyperparameter screening 13
Tab. S2 | Design settings in the AI.zymes run affording K3 14
Tab. S3 | Experimental data 14
5. Supplementary Figures 15
Fig. S1 | Ai.zymes workflow. 15

Fig. S2 | MD screening of the 96 most promising designs 16
Fig. S3 | Experimental Data. 17
Fig. S4 | Alphafold3-predicted oligomerization of KSI. 18
Fig. S5 | Score fluctuations in WT and K3. 19
6. Sequences 20
6.1. Design sequences 20
6.2. wtKSI sequence 20
6.3. K3 sequence 20
6.4. Plasmid Map of K3 20
7. Input files 21
7.1. Rosetta parameter file 21
7.2. Rosetta Match constraint file 21
7.3. Example .xml file for RosettaRelax 22
7.4. Example .xml file for RosettaDesign 22
7.5. Transition State parameters for MD 24

### 1. AI.zymes

A brief description of the AI.zymes algorithm–including an overview of the code architecture and the design settings employed in this work–is described here. Upon peer-reviewed publication, a detailed manual will be made available at https://github.com/bunzela/AIzymes.

AI.zymes is a modular program that seamlessly combines computational methods for design, structure prediction, and machine learning in a coherent enzyme design workflow (Fig. S1). To that end, AI.zymes employs a controller function that orchestrates the design process. The controller collects information from the designs and stores them in a shared database, selects which variants to submit for design, and decides what type of design or structure prediction algorithms to run with the selected variant. The controller is designed to manage a user-defined number of parallel design jobs, automatically starting a new job as soon as a previous one is completed.

#### 1.1. Basic concepts

##### 1.1.1. Evolutionary "rounds" in AI.zymes.

The evolutionary algorithm in AI.zymes is not based on performing classical evolutionary rounds comprising mutagenesis and screening. Instead, the algorithm aims to maximize performance by constantly running a set number of design jobs set by `max_jobs`. Whenever the number of running jobs falls below `max_jobs`, AI.zymes spawns another design or structure prediction job. To choose the parent variant for design, AI.zymes performs multi-objective Boltzmann selection. Importantly, all designs are treated equally during Boltzmann selection, which selects from the growing pool of designs irrespective of which round the designs stemmed. Thus, design is not limited by a maximum number of rounds but a maximum number of designs (`max_designs`).

##### 1.1.2. Scoring metrics.

AI.zymes employs various scoring metrics to assess various properties relevant to enzyme activity. The `total_score` corresponds to the total energy of the system. This score reflects the `total_score` of a structure in Rosetta and is referred to as "stability score" in the main text. The `interface_score` corresponds to the binding energy of the ligand to the protein. This score was calculated using Rosetta's InterfaceScoreCalculator mover and is referred to as the "interface score" in the main text.[25] The `catalytic_score` corresponds to the score of the catalytic interaction. It is calculated from the ideal interaction geometry described in a Rosetta_Match constraint file format (see 7.2 Rosetta Match constraint file).[24] Finally, the `efield_score` describes the electric fields along the scissile C-H bond. Fields were calculated using FieldTools,[23] and this score is referred to as the "electric field" in the main text.

Note that each variant in AI.zymes can have two sets of scores. All variants have scores from their initial design run. Variants selected for redesign also undergo structure prediction and relaxation, generating an updated set of scores. If available, Boltzmann selection is based on these updated scores; otherwise, the original design scores are used.

Furthermore, to run evolution in a forward-thinking manner, where the scores of the direct descendants of a variant can be included during screening, a metric called "potential" was introduced. The potential is the average of the score of the current variant and the scores of all its direct descendants.

##### 1.2. System startup

To start AI.zymes, the system must be set up using `setup_aizymes()`, which only runs once. The `setup_aizymes()` function creates the overall file structure and stores all design-relevant parameters in the `variables.json` file. Among others, these include the name of the protein, ligand, and which residues can be designed. Furthermore, general settings can be set, such as how many jobs may run in parallel and how many designs will be made in total. Following the initial setup, `startup_controller()` reads these parameters from the `variables.json` file and initializes all other required variables. Thus, `startup_controller()` is executed every time the code is restarted.

##### 1.3. Controller

The controller is the central program of AI.zymes. It constantly cycles between the following steps:

1. `check_running_jobs()` determines how many jobs are currently running on the system. If `MAX_JOBS` are running, the controller sleeps for 20 s and rechecks the number of running jobs, or else the controller initiates a new design cycle.
2. `update_scores()` iterates through all designs and updates the ALL_SCORES.csv database. `update_scores()` also unblocks all indices for which the structure prediction runs are completed.
3. `check_parent_done()` checks if the initial non-evolutionary design for the parent variant has concluded. If fewer designs than `N_PARENT_JOBS` have been made, the controller skips the Boltzmann selection and instead runs `start_parent_design()`, starting a `run_RosettaDesign()` job on the parent structure.
4. `boltzmann_selection()` initiates the Boltzmann selection to identify the next scaffold for design. For a description of the Boltzmann selection process, see 1.4Boltzmann selection.
5. `start_calculation()` finally starts the design based on the selected input structure. For a description of the available design modules, see 1.5 Available Modules in AI.zymes.. `start_calculation()` checks if there is a structure of the selected design that went through `ESMfold_RosettaRelax()`. If not, the controller will start the structure prediction and block the selected index. If there is a relaxed structure, the controller will generate a new index into which the design will be stored. This involves creating a folder for the new design and appending the `all_scores.csv` file with the selected index. To allow for a blend of design methods, the selected design will use ProteinMPNN instead of RosettaDesign with a user-defined probability `ProteinMPNN_PROB`.

##### 1.4. Boltzmann selection.

In contrast to all other scores, the `catalytic_score` has a clear minimum, with a score of zero reflecting a perfect agreement of the design with the target catalytic geometry. Thus, catalytic_score was not included during Boltzmann selection but used as a binary cutoff to exclude variants before selection. To that end, the mean plus one standard deviation of the catalytic_score of all designs is calculated, and all variants with a catalytic_score below that cutoff are removed. Additionally, structures that are currently undergoing structure prediction are excluded from Boltzmann selection to prevent redundant structure prediction on the same structure.

Boltzmann selection is based on the design potentials and not on their scores (see 1.1.2. Scoring metrics.). Boltzmann selection is performed on the combined_potential, which is calculated from the average of the z-score standardized total_potential, interface_potential, and efield_potential (Eq. 1, Z = standardized z-score, x = raw potential, μ = average potential, σ = standard deviation). Note that during normalization, potentials for which lower values are better are inverted (total_potential and interface_potential). Thus, higher combined_potentials correspond to better variants. Boltzmann selection is performed at a user-defined temperature kbt_boltzmann. To increase selection stringency during design, kbt_boltzmann decreases with each new design from an initial KBT_BOLTZMANN value with a KBT_BOLTZMANN_DECAY rate in a single exponential decay.

$$Z = \frac{x - \mu}{\sigma} \qquad \text{Eq. 1}$$

##### 1.4.1. Database

The all_scores.csv database is the primary file containing all information from an AI.zymes run. Among other details, the database includes data on the specific settings for making each design and its resulting scores. Additionally, all_scores.csv keeps track of which structures are currently undergoing structure prediction and are therefore excluded from Boltzmann selection. This exclusion prevents redundant structure prediction from being performed multiple times on the same structure.

#### 1.5. Available Modules in AI.zymes.

Currently, AI.zymes has established run_RosettaDesign() and run_ProteinMPNN() for design. For structure prediction and scoring, run_ESMfold_RosettaRelax() and calc_efields_score() have been established. Importantly, AI.zymes is built highly modularly, facilitating the future addition of other protein engineering packages to augment the computational evolution algorithm.

##### 1.5.1. Design with RosettaDesign.

**`run_RosettaDesign()`** setups and submits a RosettaDesign run for the selected index based on RosettaScripts.[10] AI.zymes dynamically generates the input .xml files that control RosettaScripts. An example .xml file can be found in 7.4 Example .xml file for RosettaDesign. Briefly, a geometry bias from the RosettaMatch constraint file is introduced with the AddOrRemoveMatchCsts (7.2 Rosetta Match constraint file), and the protein is repacked and minimized using the EnzRepackMinimize mover. Subsequently, the protein is designed using FastDesign[6,7] for 3 repeats while applying a bias to the input sequence using the FavorSequenceProfile mover with a weight of **`CST_WEIGHT`**. Designable active-site residues are defined with **`DESIGN`**, and all mutations but cys are permitted. For the catalytic residue, only glu and asp are permitted. After design, the protein is relaxed for 1 repeat with FastRelax mover[7,27] without the geometry bias from the RosettaMatch constraint file. The final scores, including the **`interface_score`** calculated with InterfaceScoreCalculator,[25] are given for the relaxed structure.

##### 1.5.2. Design using ProteinMPNN.

**`run_ProteinMPNN()`** setups and submits a ProteinMPNN run for the selected index using its sequence as input.[9] A bias_by_res.json file is generated that applies a bias to the input sequence. The input structure was parsed using the ProteinMPNN helper scripts (parse_multiple_chains.py, assign_fixed_chains.py, make_fixed_positions_dict.py) to define the target chain and specify the fixed positions. Residues in **`DESIGN`** are excluded from design with ProteinMPNN and are only designed with Rosetta. ProteinMPNN is executed with the specified sampling temperature **`ProteinMPNN_T`** and the generated bias file, producing a set of 100 candidate sequences. The highest-scoring sequence in terms of global_score is used for the subsequent modeling steps. Because ProteinMPNN does not provide a structure but only a sequence, **`run_ProteinMPNN()`** always spawns a **`run_ESMfold_RosettaRelax()`** to generate a structure and scores for Boltzmann selection.

##### 1.5.3. Structure prediction with ESMfold and RosettaRelax.

`run_ESMfold_RosettaRelax()` using both the sequence and structure as input to predict the protein structure with ESMFold[13]. Because ESMFold[13] cannot predict the structure of the substrate-enzyme complex, the coordinates of the ligand are transferred from the parent structure file into the predicted structure after the alignment of the two structures. Afterward, sidechains are stripped from the ESMfold model, and the resulting backbone-ligand complex is repacked and relaxed using Rosetta with the FastRelax mover.[7,27] For designs that do not provide a structure (e.g., ProteinMPNN), `run_ESMfold_RosettaRelax()` uses the structure of the parent variant as input.

##### 1.5.4. Electric field calculations.

`calc_efields_score()` is used to determine active-site electric fields of the input structures. Electric field calculation is performed with *FieldTools* (https://github.com/bunzela/FieldTools).[23] Fields are calculated using point charges from the ff19SB AMBER forcefield.[35] All stated fields correspond to the effective field along the scissile C-H bond. FieldTools relies on Coulomb's law using the point charges from the system's topology file and the coordinates from the input structure or trajectory to calculate the electric field along a target bond. FieldTools calculates electric field vectors $\vec{E}$ using the Coulomb constant $k_e$ and the vector $\vec{r}_i$ from the center of the C-H bond to the charges $Q_i$ in the system (Eq. 2). Subsequently, the effective field $E_{eff}$ projected along the C-H bond was calculated from the scalar product of the directional unity vector $\vec{d}$ along that bond and the total field $\vec{E}$ (Eq. 3). To quantify electric field effects, the analysis presented here focuses on the magnitude of the field vector $E_{eff}$ (Eq. 4).

$$\vec{E} = \sum_i k_e e \frac{Q_i}{\vec{r}_i^2} \quad \text{Eq. 2}$$

$$\vec{E}_{eff} = \vec{E} \cdot \vec{d} \quad \text{Eq. 3}$$

$$E_{eff} = \left|\vec{E}_{eff}\right| \quad \text{Eq. 4}$$

#### 2. Molecular dynamics simulations

After completing the main AI.zymes run, the best 96 variants were selected for further analysis with MD simulations. To eliminate redundancy, duplicate sequences were identified, and only the variant with the highest `combined_score` was kept for each unique sequence. The final selection was based on the `combined_score` and the diversity of the residues at the active side (residues 14, 18, 38, 54, 58, 63, 65, 82, 84, 97, 101, 112, 114).

**System building.** Molecular dynamics (MD) simulations were run using AMBER22[36] with GPU acceleration and analyzed using pytraj.[37] Structures were protonated with antechamber,[38] parameterized using AMBER19SB,[35] and solvated using the TIP3P[39] water model with a minimum of 10 Å between the solute and box edge. Any net charge was neutralized with uniform neutralizing plasma within AMBER. The geometry of the chemical transition state of 5-nitrobenzisoxazole with acetate was determined using DFT (B3LYP/6-311+G(d,p), SCRF=water, Gaussian16).[40,41] Ligand charges were generated by the RED server[42], which uses RESP[34] fitting. The parameters for the TS model are given in 7.5. Transition State parameters for MD.

**MD simulations.** Each input structure was minimized and heated from 0.1 to 298 K over 0.05 ns. During heating, the XYZ coordinates of the Cα atoms, catalytic base, and ligand coordinates were restrained with a weight of 100 kcal mol $Å^{-2}$). These restraints were relaxed over 0.5 ns (restraint weights 20, 8, 4, 2, 1 kcal mol $Å^{-2}$ for 0.1 ns each), and the structures were equilibrated for 2 ns with no positional restraints in the NVT ensemble. Apo structures were treated similarly to the holo-structures but without the ligand restraint. Production MD was run in the NPT ensemble using the Berendsen Barostat with a coupling constant of 1 ps and Langevin dynamics for temperature control with a collision frequency of 1 $ps^{-1}$ at 298 K for 10 ns.

**MD analysis.** Five features were calculated from MD simulation data to choose the final variants for experimental testing. Each feature was quantified by the average calculated over the final 2 ns of the 10 ns production run (0.1 ns/snapshot, 20 snapshots) (Fig. S2). Cutoff values were chosen according to the observed distributions to select only the top 96 variants.

1. The root mean squared deviation (RMSD) of the chemical transition state from the initial structure was calculated to assess the stability of the ligand at the active site. Variants in which the all-atom ligand RMSD was greater than 1.3 Å were discarded.
2. The distance of the chemical transition state to the catalytic base (base Cγ to the partially deprotonated C on the ligand) was assessed to ensure a reactive ligand-base distance. Variants with an average distance greater than 3.56 Å were discarded.
3. The average electric field along the scissile CH bond was calculated because strong electric fields can be expected to improve reactivity. Variants with an average electric field of less than 175 MV/cm were discarded.
4. The average all-atom RMSD between the apo- and holo-enzyme was determined to assess the preorganization of the apo state and ensure that the structure remains in an active conformation in the absence of the ligand. The apo-holo RMSD change was only calculated for residues that were designable. Variants with an average apo-holo RMSD greater than 0.9 Å were discarded.
5. The cumulative sum of the oxygen radial distribution function of water molecules around the catalytic base up to 4.5 Å was determined to identify structures in which the catalytic base is desolvated and, therefore, more reactive. Variants with an average sum greater than 0.01 were discarded.

Of the 96 variants submitted for MD simulation, 23 passed all filters. After visual inspection of the active-site geometry, 7 variants were chosen for experimental testing.

#### 3. Experimental Methods

##### 3.1. Protein production and purification

All genes were commercially synthesized by *Twist* into the PET21(+) vector. T7express cells were transformed with the vectors and plated on LB agar plates (100 µg/ml carbenicillin). A single colony was used to inoculate an overnight culture of LB medium containing 100 µg/ml carbenicillin. 1 l of LB medium containing 100 µg/ml carbenicillin was inoculated with the overnight culture in a ratio of 1:1000 and grown to an OD of 0.7 at 37 °C. Protein production was induced with 1 mM IPTG, and cells were incubated at 18 °C overnight. The cells were harvested, and the pellet was frozen at -80 °C in two aliquots of 40 ml Ni-NTA buffer A (50 mM sodium phosphate, 300 mM sodium chloride, 10 mM imidazole, pH 8.0) supplemented with 0.5 mg/ml lysozyme and 0.05 mg/ml DNAseI. After sonication on ice (1 s on, 1 s off, 60% intensity, 5 min, *Telesonic Ultrasonics*), cell debris were removed by centrifugation (18.000 rpm, 20 min, 4 °C). The soluble fraction was loaded onto a 5 ml Ni-NTA HisTrap columns (*Cytiva*), washed with 10 mM imidazole for 20 CV, and eluted with 300 mM imidazole, each in 50 mM sodium phosphate, 300 mM sodium chloride, pH 8. The buffer was exchanged over a G25 column (*Cytiva*) to 50 mM sodium phosphate, 100 mM NaCl, pH 6.81. Protein concentrations were determined by measuring the absorbance at 280 nm using their calculated extinction coefficients.[43] The following extinction coefficients were determined: ε(WT) = 5,960 $M^{-1}cm^{-1}$; ε(K3) = 20,970 $M^{-}cm^{-1}$

##### 3.2. Enzyme kinetics

Conversion of 5-nitrobenzisoxazole (*Sigma*) was assayed at 25 °C in 50 mM phosphate buffer, 100 mM NaCl containing 10% methanol, pH 7.0 (pH value after the addition of 10% methanol). Product formation was monitored spectroscopically at 380 nm (Δε = 15,800 $M^{-1}$ $cm^{-1}$). Substrate concentrations were determined by measuring the absorbance difference of the substrate added to either 10% methanol in water or 10% methanol in 1 M NaOH. Enzyme kinetics were determined at 2 µM enzyme concentration $E_0$. $k_{cat}$ and $K_M$ values were determined by fitting the initial velocities $v_0$ between 250 and 1500 µM substrate concentration to the Michaelis–Menten equation (Eq. 5). All $v_0$ values were corrected by the background reaction of 5-nitrobenzisoxazole in the absence of enzyme. pH-Rate profiles were determined by mixing 20 µl enzyme (20 mM phosphate, 20 mM NaCl, pH 8.0) with 160 µl buffer (50 mM phosphate, 50 mM acetate, 50 mM CHES, 100 mM NaCl, pH between 4 and 10) and 20 µl substrate in methanol (50 µM 5-nitrobenzisoxazole, final concentration). p$K_a$ values were determined by fitting the pH-rate profile to Eq. 6. All pH values reported in this paper correspond to the final pH of the solution after the addition of methanol.

$$\mathrm{v_0/E_0} = \frac{k_{\mathrm{cat}} \cdot [S]}{K_{\mathrm{m}} + [S]} \qquad \text{Eq. 5}$$

$$\mathrm{v_0/E_0} = \frac{(\mathrm{v_0/E_0})_{\mathrm{max}}}{1 + 10^{\mathrm{pH-p}K_{\mathrm{a}}}} \qquad \text{Eq. 6}$$

##### 3.3. Circular dichroism

Circular dichroism data were recorded with an Applied Photophysics Chirascan Plus circular dichroism spectropolarimeter. 1-4 µM Enzymes were assayed in 20 mM phosphate 20 mM NaCl, pH 8.0. The mean residual ellipticity (MRE) was calculated from the measured ellipticity Θ using a pathlength of 1 mm (d) and 139 residues (n, Eq. 7). Melting curves were recorded at 218 nm. The melting temperature $T_m$ of the WT was determined using Eq. 8 and the fitting parameters a, b, and c.

$$\mathrm{MRE} = \frac{\Theta}{\mathrm{d \cdot c \cdot (n-1)}} \qquad \text{Eq. 7}$$

$$\mathrm{MRE(T)} = \mathrm{a} + \frac{\mathrm{b}}{1 + 10^{\mathrm{c \cdot (T - T_m)}}} \qquad \text{Eq. 8}$$

### 4. Supplementary Tables

#### Tab. S1 | Hyperparameter screening

| CST_ WEIGHT | KBT_ BOLTZMANN | KBT_ BOLTZMANN_DECAY | Protein MPNN_ PROB | PMPNN_BIAS | best combined_score | designs |
|---|---|---|---|---|---|---|
| **1** | 0.08 | - | - | - | 0.66 | 4000 |
| **2** | 0.08 | - | - | - | 0.61 | 4000 |
| **3** | 0.08 | - | - | - | 0.54 | 4000 |
| **4** | 0.08 | - | - | - | 0.47 | 4000 |
| **1** | 0.5 | 0.001 | - | - | 0.67 | 4000 |
| **1** | 0.5 | 0.0005 | - | - | 0.67 | 4000 |
| **1** | 0.5 | 0.0003 | - | - | 0.68 | 4000 |
| **1** | 0.5 | 0.0003 | 0.3 | - | 0.76 | 4000 |
| **1** [a] | 0.5 [a] | 0.0003 [a] | 0.3 [a] | 0.5 [a] | 0.79 [a] | 4000 [a] |
| **1** | 0.5 | 0.0003 | 0.3 | 1.0 | 0.73 | 4000 |
| **1** | 0.5 | 0.0003 | 0.3 | 50 | 0.65 | 4000 |

[a] The final settings to create K3 are highlighted in blue.

**Tab. S2 | Design settings in the AI.zymes run affording K3**

| parameter | value |
|---|---|
| DESIGN | 7,10,11,14,15,18,26,29,30,38,54,55,58, 59,63,65,71,73,78,80,82,84,86,93,95, 97,99,101,103,109,112,114,116,121 |
| MAX_JOBS | 250 |
| N_PARENT_JOBS | 200 |
| MAX_DESIGNS | 10000 |
| KBT_BOLTZMANN [a] | 0.5 |
| KBT_BOLTZMANN_DECAY [a] | 0.0003 |
| CST_WEIGHT | 1 |
| ProteinMPNN_PROB | 0.3 |
| ProteinMPNN_BIAS | 0.5 |
| ProteinMPNN_T | 0.1 |

[a] In AI.zymes, KBT_BOLTZMANN is supplied as a list, of which the first value is the initial KBT value and the second one the decay rate constant.

**Tab. S3 | Experimental data**

| Variant | $v_0/E_0$ ($s^{-1}$) [a,b] | relative activity [a,b] | mutations |
|---|---|---|---|
| **WT** | 0.24 ± 0.02 | 1.0 | 0 |
| **K1** | inactive | inactive | 73 |
| **K2** | 0.09 ± 0.02 | 0.4 ± 0.1 | 72 |
| **K3** | 0.64 ± 0.07 | 2.7 ± 0.4 | 63 |
| **K4** | 0.023 ± 0.03 | 0.1 ± 0.1 | 77 |
| **K5** | 0.015 ± 0.001 | 0.1 ± 0.1 | 71 |
| **K6** | 0.012 ± 0.008 | 0.1 ± 0.1 | 75 |
| **K7** | 0.27 ± 0.02 | 1.1 ± 0.1 | 63 |

[a] $v_0/E_0$ values were determined at 1.5 mM 5-nitrobenzisoxazole at pH 8.0.
[b] Errors correspond to the standard deviation of 4 technical replicates.

#### 5. Supplementary Figures

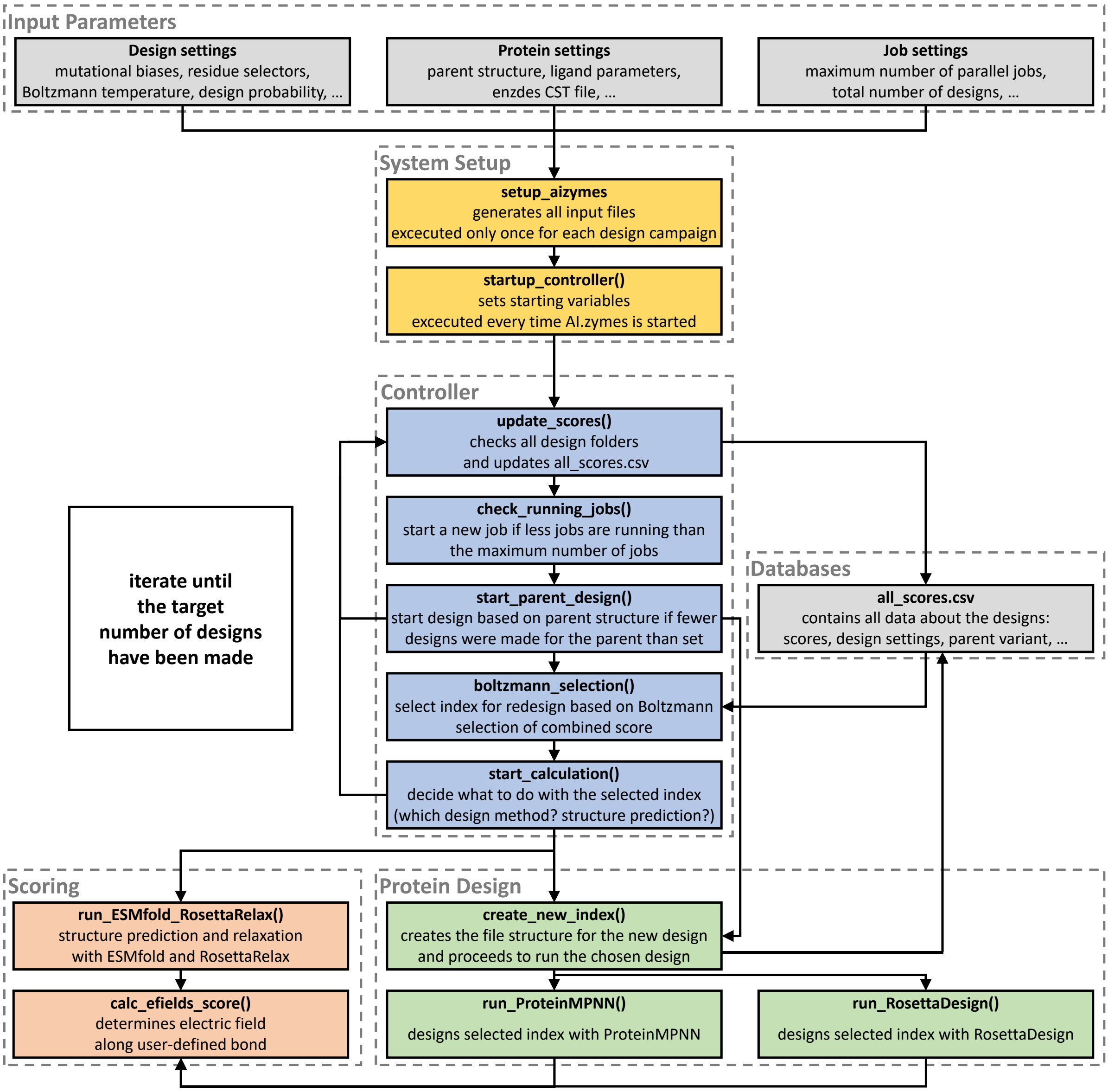


**Fig. S1 | Ai.zymes workflow.**

Based on a range of input parameters specifying the design settings, the target protein, and the overall design jobs (grey), AI.zymes is set up, and the controller is started (yellow). The controller (blue) is the core function of AI.zymes and controls all aspects of the design and screening cycle. The controller either submits variants for design (green) or structural analysis and scoring (red). All information on the designs is collected in all_scores.csv (grey). A detailed description of the AI.zymes workflow can be found in 1. AI.zymes.

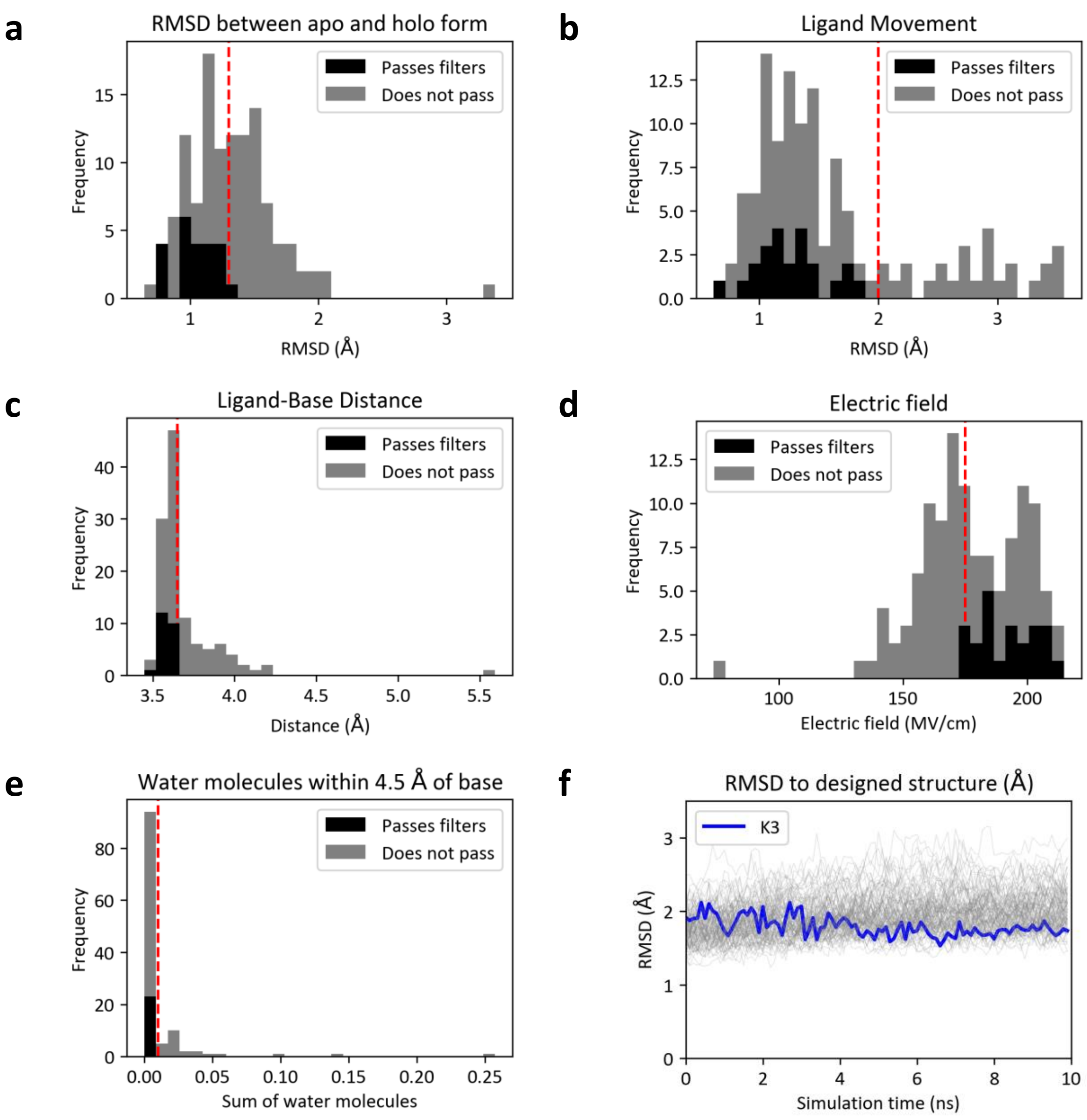


**Fig. S2 | MD screening of the 96 most promising designs**

**a-e)** Histograms showing the distribution of features used to select variants for experimental testing as described in 2. Molecular dynamics simulations. Grey bars indicate all variants screened using MD; black bars indicate the variants that pass all filters. Cut-offs for each feature are shown in red. All features are calculated from the last 2 ns of a 10 ns MD simulation. **f)** RMSD plots for the 96 most promising variants (RMSD calculated relative to the design structure before MD). K3 is shown in blue.

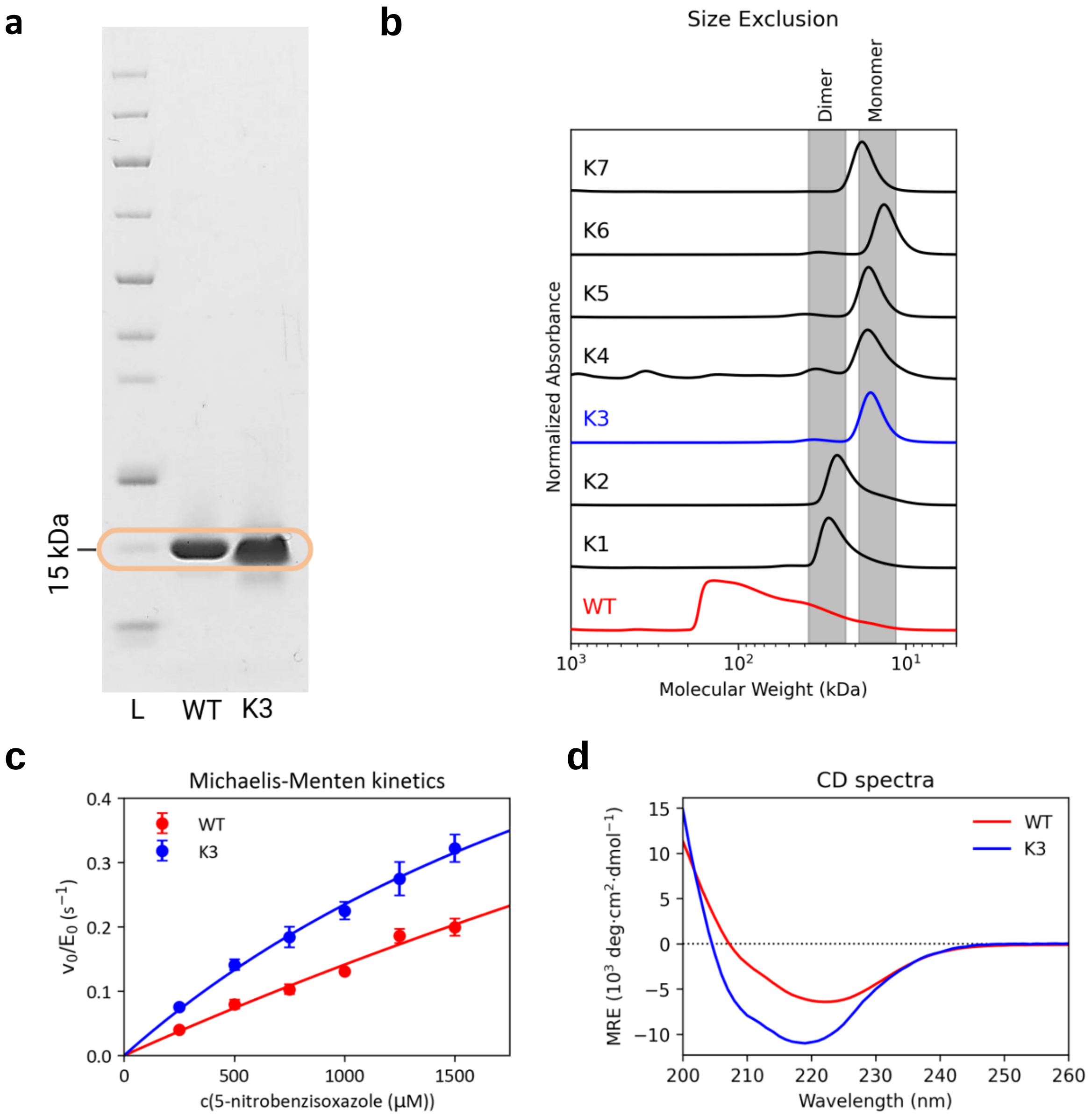


**Fig. S3 | Experimental Data.**

**a)** The WT and K3 KSI variants were obtained in high purity after Ni-NTA and buffer exchange chromatography. **b)** Analytical size exclusion chromatography revealed that K3 to K7 are present as monomers, whereas K2 and K1 form dimers. WT KSI forms a mixture of higher oligomers. **c)** Michaelis Menten kinetics of WT ($k_{cat}$ = 1.0 ± 0.03 $s^{-1}$, $k_{cat}/K_m$ = 150 ± 10 $M^{-1}s^{-1}$) and K3 ($k_{cat}$ = 4.6 ± 2.6 $s^{-1}$, $k_{cat}/K_m$ = 310 ± 10 $M^{-1}s^{-1}$) at pH 7.0 . **d)** CD spectra of WT and K3 at 25°C.

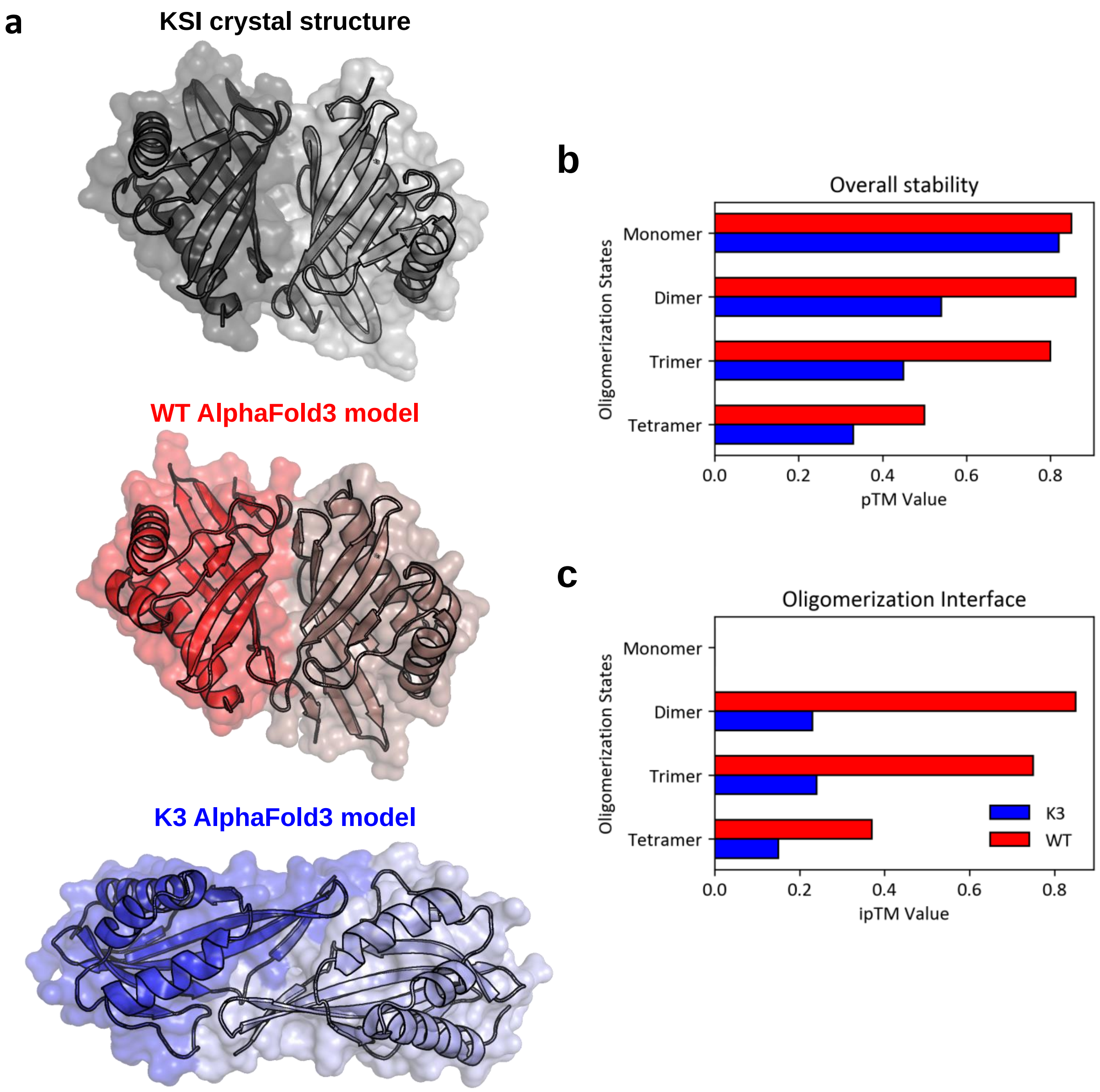


**Fig. S4 | Alphafold3-predicted oligomerization of KSI.**

**a)** Overlay of the WT dimeric crystal structure (PDB: 1OHP, light blue)[26] and the AF3 predicted structure of the WT dimer (red). **b)** The AF3 predicted dimer of K3 has a low-confidence and non-native dimer interface. **c)** Structural accuracy (pTM; left) and interface accuracy (ipTM; right) values predicted by AF3 for various oligomerization states of D38N tKSI (WT), K3 and L6. TM values below 0.5 are considered unlikely folds or interfaces.

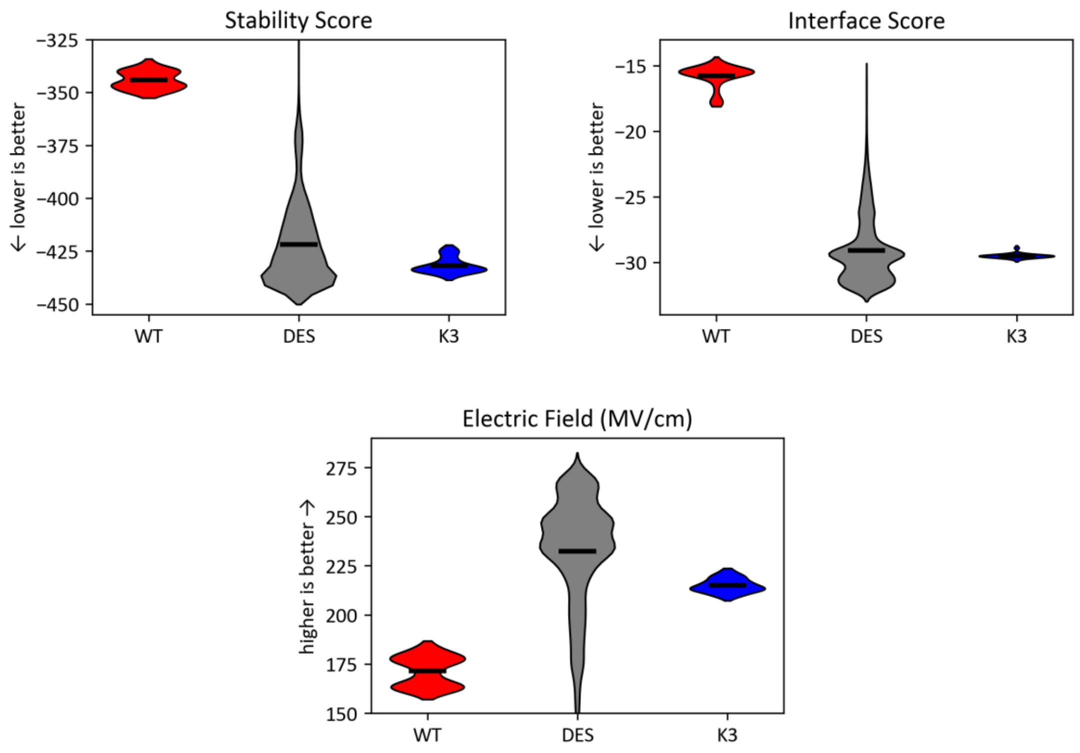


**WT - ESMfold & RosettaRelaxed, 100 structures**
**DES - Design Run, 10.000 structures**
**K3 - ESMfold & RosettaRelaxed, 100 structures**

**Fig. S5 | Score fluctuations in WT and K3.**

To assess the score fluctuations within WT (red) and K3 (blue), each structure was submitted 100 times to ESMfold and RosettaRelax. Because RosettaRelax is non-deterministic, the resulting variants have varying scores. The stability score, interface score, and electric field are significantly different in WT and K3. For comparison, the scores from all 10,000 designs created during the AI.zymes run are also shown (grey).

### 6. Sequences

#### 6.1. Design sequences

```
wtKSI   MHHHHHHHENL YFQGMNTPEH MTAVVQRYVA ALNAGDLDGI VALFADDATV ENPVGSEPRS GTAAIREFYA
K1      .......... .....C.... IK...EEWIK LV..R..E.L .....E..VL DI.P....IR .RE...AWLT
K2      .......... ....A..... IRQ..LDWFR YV..R....L ........VL DV....T.KV .KE..AAWI.
K3      .......... .....L...Q I..L..QLM. ....K....L ...W.E..VL .V......HR .RE...AW.T
K4      .......... .....L.D.Q IK.L.LELME K...K....L .....E.GVL .V.A.AP.HV .RE...A.LT
K5      .......... ....AC.... IK...REWLE YV..R....L ........VL DI.P....IR .RE...AWLT
K6      .......... ....AC.... IK...RKWFE YV..R....L ........V. HV.P.A..IV .RE...AWLT
K7      .......... .........N I....NTV.. .I..K.I..L ...W....VL .I.A....HK .PD...AWLT

wtKSI   NSLKLPLAVE LTQEVRAVAN EAAFAFIVSF EYQGRKTVVA PIDHFRFNGA GKVVSMRALF GEKNIHAGA
K1      EW....FT.K IITPIEVDG. TVT.DVEIT. T.N.K.VT.R RR.VLT..EE ..I.H..WE. SPDD.EV.E
K2      EW..F.FTIR .VTPIEIDG. TVT.D.EIT. TLN.K.V..K RR.VWT..EE ..I.RLD.H. SLDD.TV..
K3      EF..RDYKF. .VEPIQVEG. .VS.R.TL.W .EN...V..D IT..IT..E. ..I.RLV.D. S.AD.TVLE
K4      EF..TDYKF. .KEPIEVEG. S.T.N.DLT. .EN.KLVIID IT..IE..DE ..ITRLV.D. T.SD.KYIE
K5      EW....FTIK IITPIEVDG. TVT.DVEIT. T.N.K.V..R RR.VWT..EE ..I.H..WE. S.DD.EV.E
K6      EW..KDFTLK .ISPIEVDG. TVT.DVEIT. TKD.K.V..H RR.VWT..EE ..I.E.EWE. SPDD.EV.E
K7      EW....YQ.. ..TPLDIEG. KVT.D.V.T. .KD.KLF..R RT..WT..DE ..I.RLEV.. S.DD.TVLE
```

#### 6.2. wtKSI sequence

MHHHHHHHENL YFQGMNTPEH MTAVVQRYVA ALNAGDLDGI VALFADDATV ENPVGSEPRS GTAAIREFYA NSLKLPLAVE
LTQEVRAVAN EAAFAFIVSF EYQGRKTVVA PIDHFRFNGA GKVVSMRALF GEKNIHAGA

TTAACTTTAA GAAGGAGATA TACATATGCA CCATCATCAT CATCACGAAA ATTTGTACTT CCAGGGAATG AATACCCCAG
AACACATGAC AGCAGTAGTC CAGCGCTATG TTGCGGCATT AAACGCCGGT GATCTGGACG GCATAGTTGC CCTGTTTGCT
GACGACGCCA CTGTTGAAAA TCCAGTGGGA AGCGAGCCTC GTTCTGGTAC GGCTGCGATT CGTGAATTCT ATGCAAACTC
ACTGAAACTG CCTTTAGCGG TCGAACTGAC ACAGGAAGTA CGTGCCGTCG CCAATGAAGC CGCTTTTGCC TTTATAGTTT
CTTTTGAGTA CCAAGGACGC AAAACAGTAG TTGCTCCGAT AGACCATTTT CGTTTTAATG GAGCAGGGAA AGTGGTTTCG
ATGCGCGCCC TTTTTGGCGA AAAAAATATT CACGCTGGCG CCTAATGA

#### 6.3. K3 sequence

MHHHHHHHENL YFQGMLTPEQ ITALVQQLMA ALNAKDLDGL VALWAEDAVL EVPVGSEPHR GREAIRAWYT EFLKRDYKFE
LVEPIQVEGN EVSFRFTLWE ENGRKVVVDI TDHITFNEAG KIVRLVADFS EADITVLE

TTAACTTTAA GAAGGAGATA TACATATGCA CCATCATCAT CATCACGAAA ATTTGTACTT CCAGGGAATG CTCACTCCCG
AGCAAATTAC AGCGTTAGTG CAACAGCTGA TGGCAGCCCT CAATGCCAAG GATTTAGACG GATTAGTGGC CCTTTGGGCC
GAGGATGCCG TTCTGGAAGT ACCGGTCGGA TCAGAACCTC ATCGGGGACG AGAAGCGATA CGTGCTTGGT ATACGGAATT
CTTGAAACGT GATTATAAAT TCGAATTGGT CGAACCGATC CAGGTCGAAG GGAACGAAGT ATCTTTTCGC TTTACACTCT
CCTGGGAAGA AAACGGACGG AAAGTTGTTG TCGACATCAC AGATCACATC ACTTTTAATG AAGCAGGGAA AATTGTCAGA
CTGGTAGCAG ATTTCTCGGA AGCCGATATT ACTGTCCTTG AATGA

#### 6.4. Plasmid Map of K3

TTAACTTTAAGAAGGAGATATACATATGCACCATCATCATCATCACGAAAATTTGTACTTCCAGGGAATGCTCACTCCCGAGCAAATTACAGCGTTAGTGCAACAGCTGATGGCAGCCCTCAATGCCAAGGATTTAGACGGATTAGTGGCCCTTTGGGCCGAGGAT
GCCGTTCTGGAAGTACCGGTCGGATCAGAACCTCATCGGGGACGAGAAGCGATACGTGCTTGGTATACGGAATTCTTGAAACGTGATTATAAATTCGAATTGGTCGAACCGATCCAGGTCGAAGGGAACGAAGTATCTTTTCGCTTTACACTCTCCTGGGAAGAAA
ACGGACGGAAAGTTGTTGTCGACATCACAGATCACATCACTTTTAATGAAGCAGGGAAAATTGTCAGACTGGTAGCAGATTTCTCGGAAGCCGATATTACTGTCCTTGAATGAGCGGCCGCACTCGAGCACCACCACCACCACCACTGAGATCCGGCTGCTAACAA
AGCCCGAAAGGAAGCTGAGTTGGCTGCTGCCACCGCTGAGCAATAACTAGCATAACCCCTTGGGGCCTCTAAACGGGTCTTGAGGGGTTTTTTGCTGAAAGGAGGAACTATATCCGGATTGGCGAATGGGACGCGCCCTGTAGCGGCGCATTAAGCGCGGCGGGTG
TGGTGGTTACGCGCAGCGTGACCGCTACACTTGCCAGCGCCCTAGCGCCCGCTCCTTTCGCTTTCTTCCCTTCCTTTCTCGCCACGTTCGCCGGCTTTCCCCGTCAAGCTCTAAATCGGGGGCTCCCTTTAGGGTTCCGATTTAGTGCTTTACGGCACCTCGACCC
CAAAAAACTTGATTAGGGTGATGGTTCACGTAGTGGGCCATCGCCCTGATAGACGGTTTTTCGCCCTTTGACGTTGGAGTCCACGTTCTTTAATAGTGGACTCTTGTTCCAAACTGGAACAACACTCAACCCTATCTCGGTCTATTCTTTTGATTTATAAGGGATT
TTGCCGATTTCGGCCTATTGGTTAAAAAATGAGCTGATTTAACAAAAATTTAACGCGAATTTTAACAAAATATTAACGCTTACAATTTAGGTGGCACTTTTCGGGGAAATGTGCGCGGAACCCCTATTTGTTTATTTTTCTAAATACATTCAAATATGTATCCGCT
CATGAGACAATAACCCTGATAAATGCTTCAATAATATTGAAAAAGGAAGAGTATGAGTATTCAACATTTCCGTGTCGCCCTTATTCCCTTTTTTGCGGCATTTTGCCTTCCTGTTTTTGCTCACCCAGAAACGCTGGTGAAAGTAAAAGATGCTGAAGATCAGTTG
GGTGCACGAGTGGGTTACATCGAACTGGATCTCAACAGCGGTAAGATCCTTGAGAGTTTTCGCCCCGAAGAACGTTTTCCAATGATGAGCACTTTTAAAGTTCTGCTATGTGGCGCGGTATTATCCCGTATTGACGCCGGGCAAGAGCAACTCGGTCGCCGCATAC
ACTATTCTCAGAATGACTTGGTTGAGTACTCACCAGTCACAGAAAAGCATCTTACGGATGGCATGACAGTAAGAGAATTATGCAGTGCTGCCATAACCATGAGTGATAACACTGCGGCCAACTTACTTCTGACAACGATCGGAGGACCGAAGGAGCTAACCGCTTT
TTTGCACAACATGGGGGATCATGTAACTCGCCTTGATCGTTGGGAACCGGAGCTGAATGAAGCCATACCAAACGACGAGCGTGACACCACGATGCCTGCAGCAATGGCAACAACGTTGCGCAAACTATTAACTGGCGAACTACTTACTCTAGCTTCCCGGCAACAA
TTAATAGACTGGATGGAGGCGGATAAAGTTGCAGGACCACTTCTGCGCTCGGCCCTTCCGGCTGGCTGGTTTATTGCTGATAAATCTGGAGCCGGTGAGCGTGGGTCTCGCGGTATCATTGCAGCACTGGGGCCAGATGGTAAGCCCTCCCGTATCGTAGTTATCT
ACACGACGGGGAGTCAGGCAACTATGGATGAACGAAATAGACAGATCGCTGAGATAGGTGCCTCACTGATTAAGCATTGGTAACTGTCAGACCAAGTTTACTCATATATACTTTAGATTGATTTAAAACTTCATTTTTAATTTAAAAGGATCTAGGTGAAGATCCT
TTTTGATAATCTCATGACCAAAATCCCTTAACGTGAGTTTTCGTTCCACTGAGCGTCAGACCCCGTAGAAAAGATCAAAGGATCTTCTTGAGATCCTTTTTTTCTGCGCGTAATCTGCTGCTTGCAAACAAAAAAACCACCGCTACCAGCGGTGGTTTGTTTGCCG
GATCAAGAGCTACCAACTCTTTTTCCGAAGGTAACTGGCTTCAGCAGAGCGCAGATACCAAATACTGTCCTTCTAGTGTAGCCGTAGTTAGGCCACCACTTCAAGAACTCTGTAGCACCGCCTACATACCTCGCTCTGCTAATCCTGTTACCAGTGGCTGCTGCCA
GTGGCGATAAGTCGTGTCTTACCGGGTTGGACTCAAGACGATAGTTACCGGATAAGGCGCAGCGGTCGGGCTGAACGGGGGGTTCGTGCACACAGCCCAGCTTGGAGCGAACGACCTACACCGAACTGAGATACCTACAGCGTGAGCTATGAGAAAGCGCCACGCT
TCCCGAAGGGAGAAAGGCGGACAGGTATCCGGTAAGCGGCAGGGTCGGAACAGGAGAGCGCACGAGGGAGCTTCCAGGGGGAAACGCCTGGTATCTTTATAGTCCTGTCGGGTTTCGCCACCTCTGACTTGAGCGTCGATTTTTGTGATGCTCGTCAGGGGGGCGG
AGCCTATGGAAAAACGCCAGCAACGCGGCCTTTTTACGGTTCCTGGCCTTTTGCTGGCCTTTTGCTCACATGTTCTTTCCTGCGTTATCCCCTGATTCTGTGGATAACCGTATTACCGCCTTTGAGTGAGCTGATACCGCTCGCCGCAGCCGAACGACCGAGCGCA
GCGAGTCAGTGAGCGAGGAAGCGGAAGAGCGCCTGATGCGGTATTTTCTCCTTACGCATCTGTGCGGTATTTCACACCGCAATGGTGCACTCTCAGTACAATCTGCTCTGATGCCGCATAGTTAAGCCAGTATACACTCCGCTATCGCTACGTGACTGGGTCATGG
CTGCGCCCCGACACCCGCCAACACCCGCTGACGCGCCCTGACGGGCTTGTCTGCTCCCGGCATCCGCTTACAGACAAGCTGTGACCGTCTCCGGGAGCTGCATGTGTCAGAGGTTTTCACCGTCATCACCGAAACGCGCGAGGCAGCTGCGGTAAAGCTCATCAGC
GTGGTCGTGAAGCGATTCACAGATGTCTGCCTGTTCATCCGCGTCCAGCTCGTTGAGTTTCTCCAGAAGCGTTAATGTCTGGCTTCTGATAAAGCGGGCCATGTTAAGGGCGGTTTTTTCCTGTTTGGTCACTGATGCCTCCGTGTAAGGGGGATTTCTGTTCATG
GGGGTAATGATACCGATGAAACGAGAGAGGATGCTCACGATACGGGTTACTGATGATGAACATGCCCGGTTACTGGAACGTTGTGAGGGTAAACAACTGGCGGTATGGATGCGGCGGGACCAGAGAAAAATCACTCAGGGTCAATGCCAGCGCTTCGTTAATACAG
ATGTAGGTGTTCCACAGGGTAGCCAGCAGCATCCTGCGATGCAGATCCGGAACATAATGGTGCAGGGCGCTGACTTCCGCGTTTCCAGACTTTACGAAACACGGAAACCGAAGACCATTCATGTTGTTGCTCAGGTCGCAGACGTTTTGCAGCAGCAGTCGCTTCA
CGTTCGCTCGCGTATCGGTGATTCATTCTGCTAACCAGTAAGGCAACCCCGCCAGCCTAGCCGGGTCCTCAACGACAGGAGCACGATCATGCGCACCCGTGGGGCCGCCATGCCGGCGATAATGGCCTGCTTCTCGCCGAAACGTTTGGTGGCGGGACCAGTGACG
AAGGCTTGAGCGAGGGCGTGCAAGATTCCGAATACCGCAAGCGACAGGCCGATCATCGTCGCGCTCCAGCGAAAGCGGTCCTCGCCGAAAATGACCCAGAGCGCTGCCGGCACCTGTCCTACGAGTTGCATGATAAAGAAGACAGTCATAAGTGCGGCGACGATAG
TCATGCCCCGCGCCCACCGGAAGGAGCTGACTGGGTTGAAGGCTCTCAAGGGCATCGGTCGAGATCCCGGTGCCTAATGAGTGAGCTAACTTACATTAATTGCGTTGCGCTCACTGCCCGCTTTCCAGTCGGGAAACCTGTCGTGCCAGCTGCATTAATGAATCGG
CCAACGCGCGGGGAGAGGCGGTTTGCGTATTGGGCGCCAGGGTGGTTTTTCTTTTCACCAGTGAGACGGGCAACAGCTGATTGCCCTTCACCGCCTGGCCCTGAGAGAGTTGCAGCAAGCGGTCCACGCTGGTTTGCCCCAGCAGGCGAAAATCCTGTTTGATGGT
GGTTAACGGCGGGATATAACATGAGCTGTCTTCGGTATCGTCGTATCCCACTACCGAGATATCCGCACCAACGCGCAGCCCGGACTCGGTAATGGCGCGCATTGCGCCCAGCGCCATCTGATCGTTGGCAACCAGCATCGCAGTGGGAACGATGCCCTCATTCAGC
ATTTGCATGGTTTGTTGAAAACCGGACATGGCACTCCAGTCGCCTTCCCGTTCCGCTATCGGCTGAATTTGATTGCGAGTGAGATATTTATGCCAGCCAGCCAGACGCAGACGCGCCGAGACAGAACTTAATGGGCCCGCTAACAGCGCGATTTGCTGGTGACCCA
ATGCGACCAGATGCTCCACGCCCAGTCGCGTACCGTCTTCATGGGAGAAAATAATACTGTTGATGGGTGTCTGGTCAGAGACATCAAGAAATAACGCCGGAACATTAGTGCAGGCAGCTTCCACAGCAATGGCATCCTGGTCATCCAGCGGATAGTTAATGATCAG
CCCACTGACGCGTTGCGCGAGAAGATTGTGCACCGCCGCTTTACAGGCTTCGACGCCGCTTCGTTCTACCATCGACACCACCACGCTGGCACCCAGTTGATCGGCGCGAGATTTAATCGCCGCGACAATTTGCGACGGCGCGTGCAGGGCCAGACTGGAGGTGGCA
ACGCCAATCAGCAACGACTGTTTGCCCGCCAGTTGTTGTGCCACGCGGTTGGGAATGTAATTCAGCTCCGCCATCGCCGCTTCCACTTTTTCCCGCGTTTTCGCAGAAACGTGGCTGGCCTGGTTCACCACGCGGGAAACGGTCTGATAAGAGACACCGGCATACT
CTGCGACATCGTATAACGTTACTGGTTTCACATTCACCACCCTGAATTGACTCTCTTCCGGGCGCTATCATGCCATACCGCGAAAGGTTTTGCGCCATTCGATGGTGTCCGGGATCTCGACGCTCTCCCTTATGCGACTCCTGCATTAGGAAGCAGCCCAGTAGTA
GGTTGAGGCCGTTGAGCACCGCCGCCGCAAGGAATGGTGCATGCAAGGAGATGGCGCCCAACAGTCCCCCGGCCACGGGGCCTGCCACCATACCCACGCCGAAACAAGCGCTCATGAGCCCGAAGTGGCGAGCCCGATCTTCCCCATCGGTGATGTCGGCGATATA
GGCGCCAGCAACCGCACCTGTGGCGCCGGTGATGCCGGCCACGATGCGTCCGGCGTAGAGGATCGAGATCTCGATCCCGCGAAATTAATACGACTCACTATAGGGGAATTGTGAGCGGATAACAATTCCCCTCTAGGATCCGAATTC

### 7. Input files

#### 7.1. Rosetta parameter file

**Filename**: 5TS.prepi

```
    0    0    2

This is a remark line
molecule.res
5TS   INT  0
CORRECT     OMIT DU   BEG
  0.0000
   1  DUMM  DU    M    0  -1  -2     0.000      .0        .0      .00000
   2  DUMM  DU    M    1   0  -1     1.449      .0        .0      .00000
   3  DUMM  DU    M    2   1   0     1.523   111.21       .0      .00000
   4  O3    o     M    3   2   1     1.540   111.208  -180.000 -0.474100
   5  N2    no    M    4   3   2     1.235    75.361  -150.344  0.672900
   6  O4    o     E    5   4   3     1.234   122.610  -131.841 -0.474100
   7  C1    ca    M    5   4   3     1.450   118.700    48.520  0.018800
   8  C6    ca    S    7   5   4     1.393   118.907  -178.844 -0.452600
   9  H01   ha    E    8   7   5     1.090   121.508    -0.126  0.431000
  10  C2    ca    M    7   5   4     1.409   118.909     1.062 -0.057300
  11  H02   ha    E   10   7   5     1.090   119.590     0.094  0.154200
  12  C3    ca    M   10   7   5     1.380   120.873  -179.895 -0.559100
  13  H03   ha    E   12  10   7     1.090   120.834   179.922  0.203100
  14  C4    ca    M   12  10   7     1.418   118.312    -0.078  0.720000
  15  C5    ca    M   14  12  10     1.426   119.334     0.162  0.078900
  16  C9    cd    M   15  14  12     1.450   106.056   179.929 -0.296900
  17  H04   h4    E   16  15  14     1.090   121.755   179.952  0.732500
  18  N1    nc    M   16  15  14     1.246   116.514    -0.045 -0.190500
  19  O5    os    M   18  16  15     1.797   101.334    -0.031 -0.506800


LOOP
   C5   C6
   O5   C4

IMPROPER
   C1   O3   N2   O4
   C6   C2   C1   N2
   C5   C1   C6  H01
   C3   C1   C2  H02
   C4   C2   C3  H03
   C3   C5   C4   O5
   C4   C6   C5   C9
   C5  H04   C9   N1

DONE
STOP
```

#### 7.2. Rosetta Match constraint file

**Filename**: 5TS_enzdes_planar_tAB100.cst

```
CST::BEGIN
  TEMPLATE::   ATOM_MAP: 1 atom_name: C9 N1 O5
  TEMPLATE::   ATOM_MAP: 1 residue3:  5TS
  TEMPLATE::   ATOM_MAP: 2 atom_type: OOC
  TEMPLATE::   ATOM_MAP: 2 residue1:  DE
  CONSTRAINT:: distanceAB:     2.6 0.0 100.0   0. 0
  CONSTRAINT::    angle_A:   120.0 0.0  50.0 360. 0
  CONSTRAINT::    angle_B:   120.0 0.0  50.0 120. 0
  CONSTRAINT::  torsion_A:     0.0 0.0  35.0 180. 0
  CONSTRAINT:: torsion_AB:     0.0 0.0  100.0 180. 0
  CONSTRAINT::  torsion_B:     0.0 0.0  35.0 180. 0
CST::END
```

##### 7.3. Example .xml file for RosettaRelax

**Note:** The xml script is dynamically generated from `run_RosettaRelax()`. It is adjusted here for readability.

```
<ROSETTASCRIPTS>

  <SCOREFXNS>
    <ScoreFunction name    = "score"                   weights = "beta_nov16" >
      <Reweight scoretype  = "atom_pair_constraint"    weight  = "1" />
      <Reweight scoretype  = "angle_constraint"        weight  = "1" />
      <Reweight scoretype  = "dihedral_constraint"     weight  = "1" />
    </ScoreFunction>
    <ScoreFunction name    = "score_final"             weights = "beta_nov16" >
      <Reweight scoretype  = "atom_pair_constraint"    weight  = "1" />
      <Reweight scoretype  = "angle_constraint"        weight  = "1" />
      <Reweight scoretype  = "dihedral_constraint"     weight  = "1" />
    </ScoreFunction>
  </SCOREFXNS>

  <MOVERS>
    <FastRelax                 name                   = "mv_relax"
                               disable_design         = "false"
                               repeats                = "1" />
    <AddOrRemoveMatchCsts      name                   = "mv_add_cst"
                               cst_instruction        = "add_new"
                               cstfile                = "5TS_enzdes_planar_tAB100.cst" />
    <InterfaceScoreCalculator  name                   = "mv_inter"
                               chains                 = "X"
                               scorefxn               = "score_final" />
  </MOVERS>

  <PROTOCOLS>
    <Add mover_name="mv_relax" />
    <Add mover_name="mv_add_cst" />
    <Add mover_name="mv_inter" />
  </PROTOCOLS>

</ROSETTASCRIPTS>
```

##### 7.4. Example .xml file for RosettaDesign

**Note:** The xml script is dynamically generated from `run_RosettaDesign()`. It is adjusted here for readability.

```
<ROSETTASCRIPTS>

  <SCOREFXNS>
    <ScoreFunction             name      = "score"                  weights = "beta_nov16" >
      <Reweight                scoretype = "atom_pair_constraint"   weight  = "1" />
      <Reweight                scoretype = "angle_constraint"       weight  = "1" />
      <Reweight                scoretype = "dihedral_constraint"    weight  = "1" />
      <Reweight                scoretype = "res_type_constraint"    weight  = "1" />
    </ScoreFunction>
    <ScoreFunction             name      = "score_unconst"          weights = "beta_nov16" >
      <Reweight                scoretype = "atom_pair_constraint"   weight  = "0" />
      <Reweight                scoretype = "dihedral_constraint"    weight  = "0" />
      <Reweight                scoretype = "angle_constraint"       weight  = 0" />
    </ScoreFunction>
    <ScoreFunction             name      = "score_final"            weights = "beta_nov16" >
      <Reweight                scoretype = "atom_pair_constraint"   weight  = "1" />
      <Reweight                scoretype = "angle_constraint"       weight  = "1" />
      <Reweight                scoretype = "dihedral_constraint"    weight  = "1" />
    </ScoreFunction>
  </SCOREFXNS>
  <RESIDUE_SELECTORS>
    <Index                     name                    = "sel_design"
                               resnums = "14,18,38,54,58,63,65,82,84,97,101,112,114"/>
    <Index                     name                    = "sel_cat_0"
                               resnums                 = "99" />
```

```
    <Not                       name                   = "sel_nothing"
                               selector               = "sel_design" />
  </RESIDUE_SELECTORS>

  <TASKOPERATIONS>
    <OperateOnResidueSubset           name          = "tsk_design"
                                      selector      = "sel_design" >
     <RestrictAbsentCanonicalAASRLT   aas           = "GPAVLIMFYWHKRQNEDST" />
    </OperateOnResidueSubset>
    <OperateOnResidueSubset           name          = "tsk_cat_0"
                                      selector      = "sel_cat_0" >
      <RestrictAbsentCanonicalAASRLT  aas           = "DE" />
    </OperateOnResidueSubset>
    <OperateOnResidueSubset           name          = "tsk_nothing"
                                      selector      = "sel_nothing" >
      <PreventRepackingRLT />
    </OperateOnResidueSubset>
  </TASKOPERATIONS>

  <FILTERS>
    <HbondsToResidue           name                   = "flt_hbonds"
                               scorefxn               = "score"
                               partners               = "1"
                               residue                = "1X"
                               backbone               = "true"
                               sidechain              = "true"
                               from_other_chains      = "true"
                               from_same_chain        = "false"
                               confidence             = "0" />
  </FILTERS>

  <MOVERS>
    <FavorSequenceProfile      name                   = "mv_native"
                               weight                 = "1.0"
                               use_native             = "true"
                               matrix                 = "IDENTITY"
                               scorefxns              = "score" />
    <EnzRepackMinimize         name                   = "mv_cst_opt"
                               scorefxn_repack        = "score"
                               scorefxn_minimize      = "score_final"
                               cst_opt                = "true"
                               task_operations = "tsk_design,tsk_nothing,tsk_cat_0" />
    <AddOrRemoveMatchCsts      name                   = "mv_add_cst"
                               cst_instruction        = "add_new"
                               cstfile = "../Input/5TS_enzdes_planar_tAB100.cst" />
    <FastDesign                name                   = "mv_design"
                               disable_design         = "false"
                               task_operations        = "tsk_design,tsk_nothing,tsk_cat_0"
                               repeats                = "1"
                               ramp_down_constraints  = "false"
                               scorefxn               = "score" />
    <FastRelax                 name                   = "mv_relax"
                               disable_design         = "true"
                               task_operations        = "tsk_design,tsk_nothing,tsk_cat_0"
                               repeats                = "1"
                               ramp_down_constraints  = "false"
                               scorefxn               = "score_unconst" />
    <InterfaceScoreCalculator  name                   = "mv_inter"
                               chains                 = "X"
                               scorefxn               = "score_final" />
  </MOVERS>

  <PROTOCOLS>
    <Add mover_name="mv_add_cst" />
    <Add mover_name="mv_cst_opt" />
    <Add mover_name="mv_native" />
    <Add mover_name="mv_design" />
    <Add mover_name="mv_relax" />
    <Add mover_name="mv_inter" />
  </PROTOCOLS>

</ROSETTASCRIPTS>
```

#### 7.5. Transition State parameters for MD

**Filename**: 5TS.prepi

```
    0    0    2

This is a remark line
molecule.res
5TS   INT  0
CORRECT     OMIT DU   BEG
  0.0000
   1  DUMM  DU    M    0  -1  -2     0.000      .0        .0      .00000
   2  DUMM  DU    M    1   0  -1     1.449      .0        .0      .00000
   3  DUMM  DU    M    2   1   0     1.523   111.21       .0      .00000
   4  O3    o     M    3   2   1     1.540   111.208  -180.000 -0.474100
   5  N2    no    M    4   3   2     1.235    75.361  -150.344  0.672900
   6  O4    o     E    5   4   3     1.234   122.610  -131.841 -0.474100
   7  C1    ca    M    5   4   3     1.450   118.700    48.520  0.018800
   8  C6    ca    S    7   5   4     1.393   118.907  -178.844 -0.452600
   9  H01   ha    E    8   7   5     1.090   121.508    -0.126  0.431000
  10  C2    ca    M    7   5   4     1.409   118.909     1.062 -0.057300
  11  H02   ha    E   10   7   5     1.090   119.590     0.094  0.154200
  12  C3    ca    M   10   7   5     1.380   120.873  -179.895 -0.559100
  13  H03   ha    E   12  10   7     1.090   120.834   179.922  0.203100
  14  C4    ca    M   12  10   7     1.418   118.312    -0.078  0.720000
  15  C5    ca    M   14  12  10     1.426   119.334     0.162  0.078900
  16  C9    cd    M   15  14  12     1.450   106.056   179.929 -0.296900
  17  H04   h4    E   16  15  14     1.090   121.755   179.952  0.732500
  18  N1    nc    M   16  15  14     1.246   116.514    -0.045 -0.190500
  19  O5    os    M   18  16  15     1.797   101.334    -0.031 -0.506800


LOOP
   C5   C6
   O5   C4

IMPROPER
   C1   O3   N2   O4
   C6   C2   C1   N2
   C5   C1   C6  H01
   C3   C1   C2  H02
   C4   C2   C3  H03
   C3   C5   C4   O5
   C4   C6   C5   C9
   C5  H04   C9   N1

DONE
STOP
```

**Filename**: 5TS.frcmod

```
Remark line goes here
MASS

BOND

ANGLE

DIHE
ca-ca-cd-h4   4    2.800       180.0002.000      same as X -c2-ca-X , penalty score=232.0
ca-ca-cd-nc   4    2.800       180.0002.000      same as X -c2-ca-X , penalty score=232.0

IMPROPER
ca-o -no-o          1.1          180.0         2.0          Using the default value
ca-ca-ca-no         1.1          180.0         2.0          Using the default value
ca-ca-ca-ha         1.1          180.0         2.0          Using general improper torsional
X- X-ca-ha, penalty score=  6.0)
ca-ca-ca-os         1.1          180.0         2.0          Using the default value
ca-ca-ca-cd         1.1          180.0         2.0          Using the default value
ca-h4-cd-nc         1.1          180.0         2.0          Using the default value

NONBON
```
